# Whole Brain Correlates of Individual Differences in Skin Conductance Responses during Human Fear Conditioning

**DOI:** 10.1101/2021.04.20.440479

**Authors:** Kevin Vinberg, Jörgen Rosén, Granit Kastrati, Fredrik Åhs

## Abstract

Understanding the neural basis for individual differences in the conditioned skin conductance response (SCR) may inform on autonomic regulation in fear-related psychopathology. Previous region-of-interest (ROI) analyses have implicated the amygdala in regulating conditioned SCR, but whole brain analyses are lacking. This study examined correlations between individual differences in conditioned SCR and neural activity throughout the whole brain by using data from a large functional magnetic resonance imaging study (*N* = 285). Results show that conditioned SCR correlates with activity in the dorsal anterior cingulate cortex/anterior midcingulate cortex, anterior insula, bilateral temporoparietal junction, right frontal operculum, bilateral dorsal premotor cortex, right superior parietal lobe and midbrain. A ROI analysis replicated a correlation between amygdala activity and conditioned SCR, but amygdala contribution to SCR was modest compared with other regions. We suggest that implicated neural regions belong to a large-scale midcingulo-insular network related to salience detection and autonomic-interoceptive processing. Altered activity within this network may underlie individual differences in conditioned SCR and autonomic aspects of psychopathology.

## 1. Introduction

Differential fear conditioning is one of the most common methods for studying fear learning in laboratory settings (e.g. Fullana et al., 2016). It refers to the pairing of an initially neutral stimulus (CS+) with an aversive unconditioned stimulus (US), whereby the CS+ consequently comes to elicit increased autonomic responses relative to an unpaired neutral stimulus (CS−). Individual differences in differential fear conditioning have been linked to pathological fear and anxiety (Duits et al., 2015; Nees, Heinrich, & Flor, 2015; Lonsdorf & Merz, 2017) and are part of contemporary clinical models of these disorders (Mineka & Oehlberg, 2008; Bouton, Mineka, & Barlow, 2001; Craske, Treanor, Conway, Zbozinek, & Vervliet, 2014). Understanding the neural basis of individual differences in differential fear conditioning may improve our understanding of autonomic regulation in pathological fear and anxiety (Lonsdorf & Merz, 2017), possibly providing us with biomarkers for pathology (e.g.MacNamara et al., 2015) and ultimately informing the development of more effective treatments (see e.g. Insel 2014). It has thus been argued that fear conditioning research should increasingly focus on understanding sources to individual differences and use large sample sizes to achieve this aim (Lonsdorf & Merz, 2017).

The most common way of measuring conditioned fear is the differential Skin Conductance Response (SCR; Lonsdorf et al., 2017), referring to changes in skin conductance induced by sympathetic activation (Dawson Schell & Fillion, 2007; Wallin et al. 1981). Previous studies considering the neural correlates of individual differences in magnitude of conditioned SCR, generally defined as the average CS+ SCR score subtracted from the average CS− SCR score, have implicated the amygdala (see e.g. Labar, Gatenby, Gore, LeDoux & Phelps, 1998; Phelps, Delgado, Nearing & LeDoux et al 2004; Dunsmoor, Prince, Murty, Kragel & Labar, 2011; Petrovic, Kalisch, Pessiglione, Singer & Dolan, 2008; MacNamara et al., 2015; Marin et al., 2019). This is in line with the general understanding of fear conditioning from animal models, where a neural circuitry centered on the amygdala has been demonstrated to be responsible for physiological and behavioral conditioning (LeDoux 2000; Davis 2000). It also connects with human lesion studies demonstrating diminished or absent conditioned SCR following amygdala damage (Labar et al. 1995; Bechara et al., 1995), as well as with human neuroimaging studies considering the neural correlates of within subject variation in conditioned SCR (Cheng, Knight, Smith, Stein & Helmstetter, 2003; Knight, Nguyen, & Bandettini, 2005; Cheng, Knight, Smith & Helmstetter, 2006).

However, these previous studies of the neural correlates of individual differences in conditioned SCR have generally been based on small sample sizes (*N* < 28) and regions of interest analysis (ROIs) targeting the amygdala. Such sample sizes fall below minimum requirements for correlation analysis in fMRI research according to guidelines (Yarkoni, 2009; Yarkoni & Braver, 2010), that recommend a sample size of at least *N* = 40 for determination of inter-individual correlations.

Furthermore, the previous focus on the amygdala excludes other regional activations potentially important for explaining individual differences in conditioned SCR. In particular, the human fear conditioning task is known to activate a large set of cortical, subcortical and brainstem areas other than the amygdala (Fullana et al., 2016). In their meta-analysis, Fullana et al. (2016) proposed that neural regions consistently activated during fear conditioning collectively constitute a large-scale neural network, centered on the dorsal anterior cingulate cortex and anterior insula, that represents autonomic-interoceptive processing in response to conditioned stimuli (Fullana et al. 2016; based on findings by e.g. Cameron, 2009; Craig, 2009; Critchley & Harrison, 2013; Medford & Critchley, 2010). Based on this proposal, one would expect individual differences in autonomic conditioned responding, such as measured by SCR (Dawson Schell & Fillion, 2007), to correlate with altered activity within this broader network.

Only two previous studies that investigated the association between individual differences in fear conditioning and neural activity have met the minimum requirements for sample size suggested by Yarkoni & Braver (2010): MacNamara et al (2015; *N* = 49) and Marin et al (2019; *N* = 60). Both these studies implicated additional neural correlates besides the amygdala. In particular, the study by Marin et al. (2019) implicated the dorsal anterior cingulate cortex and insula, two major nodes within the neural network proposed to underlie fear conditioning by Fullana et al. (2016). MacNamara et al. (2015) did not report correlations in these regions but found correlates of SCR in the supplementary motor area, another area consistently activated during human fear conditioning (Fullana et al. 2016). However, significant associations from both studies were still only reported using ROI analyses and not whole brain analysis, meaning that neural correlates in areas outside the predefined ROIs may still have gone unnoticed.

As previous studies have restricted analyses to predefined areas of the brain and/or have failed to report significant whole brain correlations, we aimed to investigate the whole brain correlations of individual differences in conditioned SCR by analyzing data from a large sample of participants undergoing fear conditioning (*N* = 285). This sample size provided sufficient statistical power to detect correlations with medium effect size and above (*r* > .251) throughout the whole brain (see the Materials and Methods section for details). The use of a whole brain approach in establishing neural correlates of individual differences in conditioned SCR has potential to identify novel brain regions influencing fear conditioning and psychopathology. Besides the whole brain analysis of conditioned SCR and neural activation, a second aim of our study was to replicate previous findings of an association between individual differences in amygdala response and SCR using an ROI approach. Finally, a third aim was to determine whether areas whose activity explained significant individual variation in conditioned SCR did so independently of each other. A comparison of the relative contributions of different brain areas to conditioned SCR has potential to inform on separate neural pathways mediating individual differences in conditioned autonomic responding. Ultimately, such findings may aid the understanding of autonomic regulation in pathological fear and anxiety.

## 2. Results

### 2.1 SCR

Subjects displayed significantly larger SCR to the CS+ relative to the CS− (*t*(284) = 23.28; *p* < .001; *d* = 1.38), indicating successful conditioning. An SCR difference score between the CS+ relative to the CS− was calculated for each participant (*M* = 0.64, *SD* = 0.47) and examining the distribution of SCR difference scores revealed substantial individual differences (see Supplementary Information 1).

### 2.2 Correlation between SCR difference scores and brain responses during fear conditioning

#### 2.2.1 Whole brain analysis

Fear conditioning related brain responses (CS+>CS−) were correlated to SCR difference scores in the dorsal anterior cingulate cortex/anterior midcingulate cortex, right anterior insula, right inferior frontal gyrus/frontal operculum, bilateral temporoparietal junction/superior temporal gyrus, right superior parietal lobe/postcentral gyrus, bilateral superior frontal gyri/dorsal premotor cortex, and a right-lateralized midbrain region in areas consistent with periaqueductal gray and reticular formation. In order to ensure the reliability of our findings and facilitate comparison with prior studies, we performed the same analysis using average square root transformed raw value SCRs instead of Z transformed SCRs. We also performed the same analysis without any participant exclusion (see the Materials and Methods section regarding participant exclusion). Both these analyses resulted in a very similar pattern of correlations (see Supplementary Information 3 and 4), showing that the results seem robust to different types of normalization of SCR and to different choices regarding participant exclusion. For a summary of results, see Figure 1 and Table 1. The effects of fear conditioning on fMRI responses (CS+ > CS−) are described in Supplementary Information 6.

**Figure 1.**
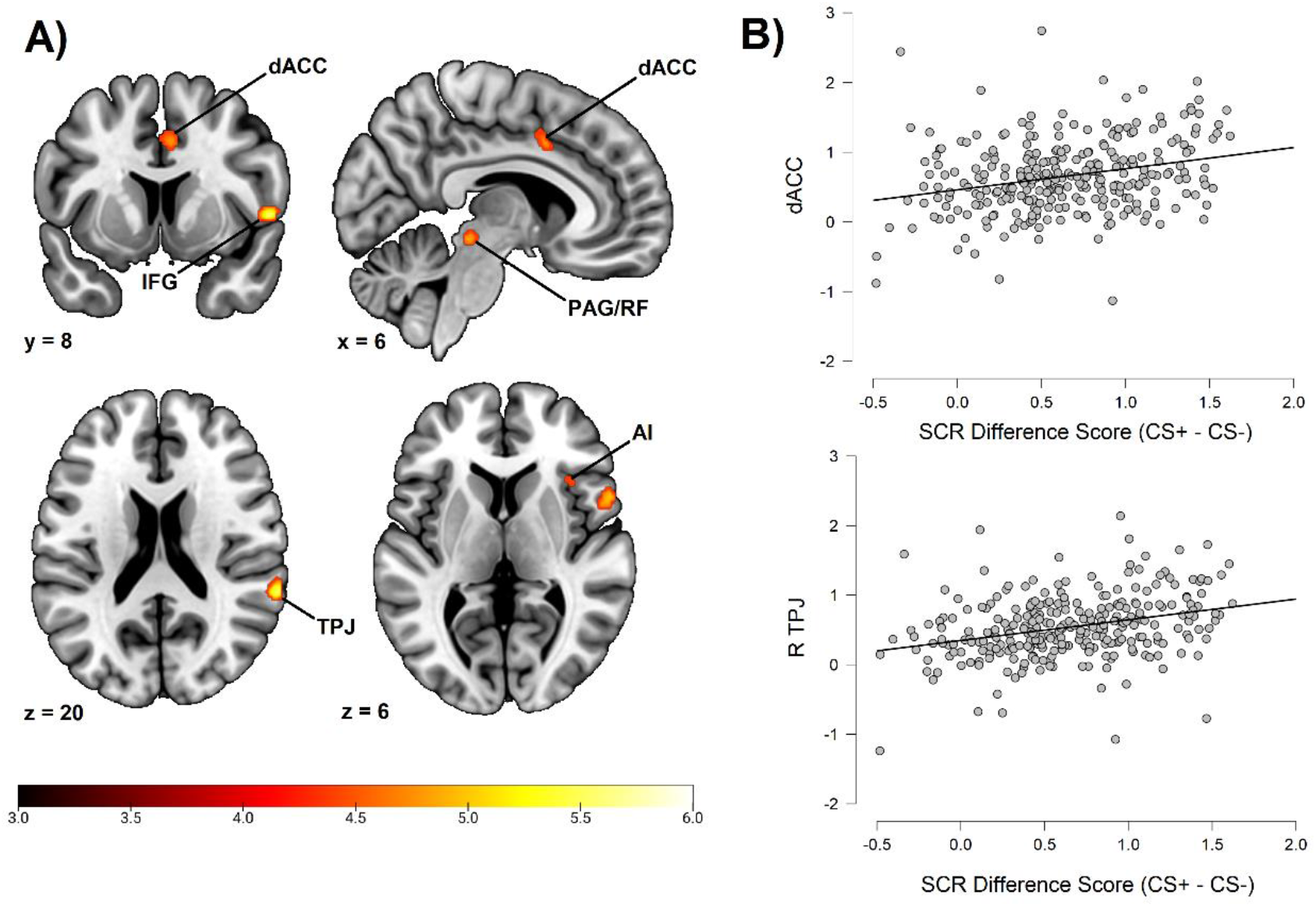
Correlation between individual differences in conditioned SCR and whole brain responses during fear conditioning. A) Activation map of key implicated neural regions. Color-coded *t* values ranges from *t* = 3 to *t* = 6. The statistical image was thresholded at *p* < 0.05 FWE-corrected and displayed on an anatomical brain template. B) Scatter plots depicting correlation between SCR difference scores and eigenvariates from significant whole brain clusters in the dorsal anterior cingulate cortex (upper panel) and the temporoparietal junction (lower panel). R = Right. dACC = dorsal Anterior Cingulate Cortex. TPJ = Temporoparietal Junction. IFG = Inferior Frontal Gyrus. PAG/SC = Periaqueductal gray/Superior Colliculus. AI = Anterior Insula.

**Table 1.** Whole Brain Correlation to Conditioned SCR

| Anatomical region | Hemisphere | Voxels | *t* | MNI Coordinates |  |  |
| --- | --- | --- | --- | --- | --- | --- |
|  |  |  |  | *x* | *y* | *z* |
| Dorsal Anterior Cingulate Cortex/Anterior Midcingulate Cortex | N/A | 50 | 4.79 | 6 | 8 | 40 |
| Anterior Insula | Right | 20 | 4.65 | 36 | 20 | 6 |
| Inferior Frontal Gyrus/Frontal Operculum | Right | 138 | 5.80 | 56 | 10 | 2 |
| Temporoparietal Junction/Superior Temporal Gyrus | Right | 81 | 5.66 | 64 | -40 | 20 |
| Superior Frontal Gyrus/Dorsal Premotor Cortex | Right | 44 | 5.21 | 18 | 0 | 68 |
| Midbrain | Right | 59 | 5.22 | 10 | -30 | -12 |
| Superior Parietal Lobe/Postcentral Gyrus | Right | 3 | 4.52 | 22 | -46 | 70 |
| Superior Frontal Gyrus/Dorsal Premotor Cortex | Left | 2 | 4.46 | -14 | -2 | 72 |
| Temporoparietal Junction/Superior Temporal Gyrus | Left | 1 | 4.37 | -62 | -36 | 22 |

*Note.* MNI coordinates and *t* values represent significant peak voxels of each cluster. Statistical significance was calculated using *t* tests implemented within the SPM software with an FWE corrected alpha level of $\alpha = .05$ .

#### 2.2.2 Correlation between individual differences in conditioned SCR and amygdala activation

While no correlation between individual differences in conditioned SCR and amygdala activation survived the statistical threshold in the whole brain analysis, a ROI analysis targeting the bilateral amygdala was performed. This analysis demonstrated significant correlations bilaterally in the amygdala (right peak MNI coordinates: 20, −2, −14; cluster size = 38 voxels; *t* = 3.68; left peak MNI coordinates: −22, −2, −16; cluster size = 8 voxels; *t* = 3.08). See figure 2.

**Figure 2.**
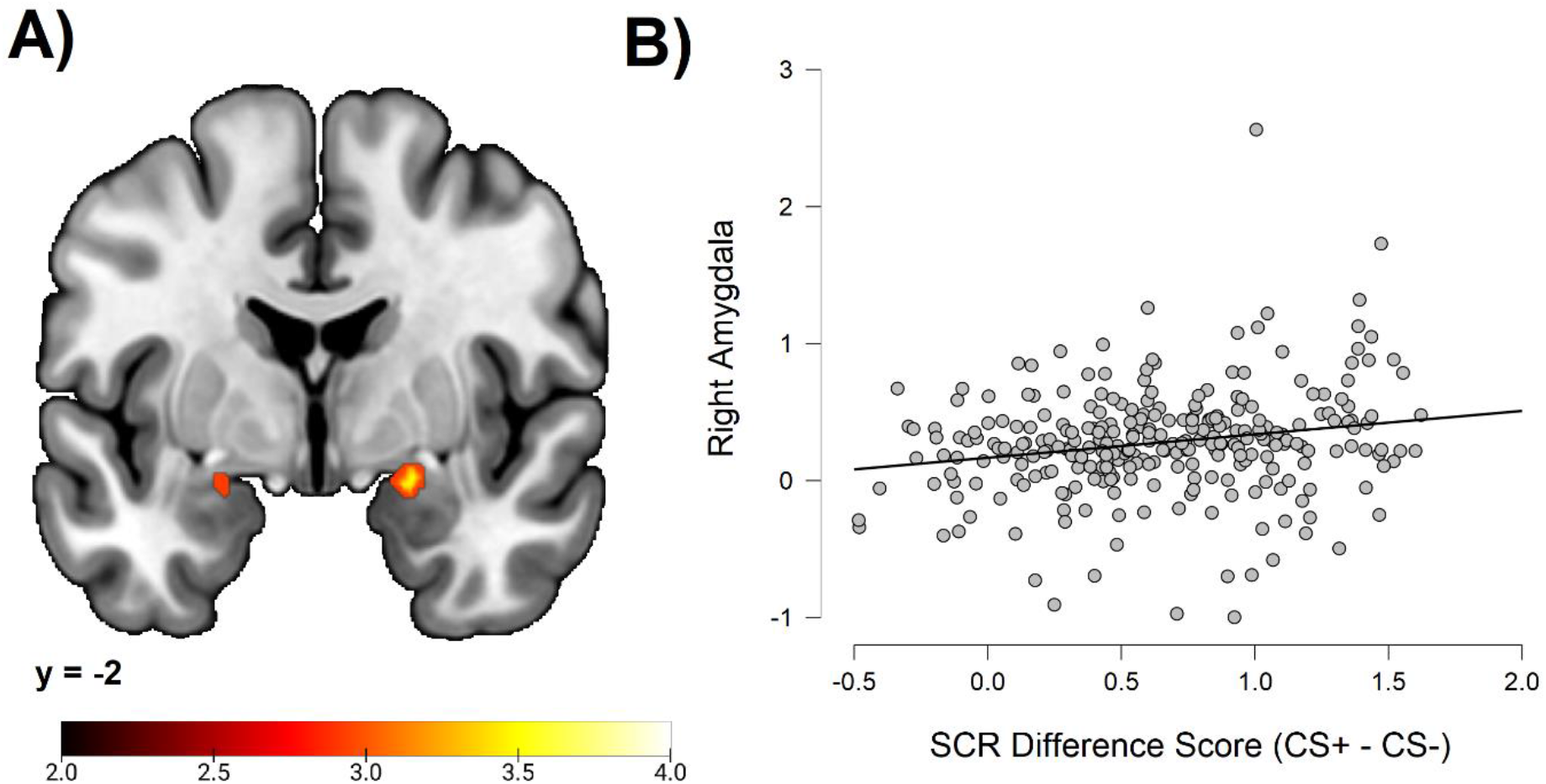
Correlations between individual differences in conditioned SCR and amygdala activation. A) Activation map depicting significant activation on coronal section at MNI Y-coordinate = −2. Color-coded *t* values ranges from *t* = 2.0 to *t* = 4.0. The statistical image was thresholded at *p* < 0.05 FWE-corrected. B) Scatter plot depicting correlation between SCR difference scores and eigenvariates from the significant right amygdala cluster within the amygdala ROI.

#### 2.2.3 Relative contribution of whole brain and amygdala correlates to individual differences in SCR

In order to examine the independent and/or shared contributions from implicated neural areas, we extracted eigenvariates from all significant clusters in the whole brain analysis and the amygdala ROI analysis and entered these as regressors in a hierarchical regression analysis. Together, regional eigenvariates demonstrated a significant correlation to conditioned SCR with a moderate effect size (*F*(1, 284) = 4.23; *r* = .38; *r*^2^ = .15; *p* < .001). By including amygdala eigenvariates in the null model and examining *r*^2^ change by adding whole brain correlates, a significant change in *r*^2^ was observed (*F*(1, 284) change = 3.69; r^2^ change = .10; *p* < .001). This meant that the whole brain correlates explained additional variance beyond that shared with the amygdala. In contrast, including regional activations from the whole brain analysis in the null model and examining *r*^2^ change by adding eigenvariates from the amygdala ROI did not result in any significant change (*F*(1,284) change = 1.50; *r*^2^ change = .01; *p* = .226). This meant that no unique contribution from amygdala activity could be demonstrated beyond that shared with other brain regions. When examining the beta weights of all regressors in the regression model (see Supplementary Information 5), we noticed significant contribution from right midbrain eigenvariates (*p* = .031). Also, when including all regional eigenvariates except the midbrain in a null model and examining *r*^2^ change by adding midbrain to the model, a significant change in *r*^2^ was observed (*F*(1,284) change = 4.70; r^2^ change = .015; *p* = .031).This meant that the midbrain demonstrated significant unique explained variance relative to all other implicated regions. Notably, however, this unique contribution could not be observed in the alternative model where the amygdala was excluded (*p* = .096; see Supplementary Information 5) indicating that this finding should be interpreted with caution since it may not replicate.

Finally, in order to determine if neural correlates to differential SCR were driven more by CS+ or CS−, we correlated eigenvariates from regions that were correlated to SCR difference scores with average raw value SCRs to the CS+ and CS− separately. Results demonstrated significant correlations between all regional BOLD eigenvariates and CS+ SCR (*p* values ≤ .001), except for right superior parietal lobe (*p* = .002), left superior frontal gyrus (*p* = .002), right amygdala (*p* = .006) and left amygdala (*p* = .029), where correlations did not survive the Bonferroni-corrected threshold (*α* = .00151; see the Materials and Methods section for details regarding the Bonferroni-correction).

However, no regional eigenvariates evidenced significant correlation to CS− SCR (*p* values > .00151) and all regional eigenvariates were significantly more correlated to CS+ SCR than CS− SCR (*p* values < .00151) except for right amygdala (*p* = .018) and left amygdala (*p* = .011). This indicated that neural correlations to differential SCR were mainly driven by increased responding to the CS+.

## 3. Discussion

Individual differences in conditioned SCR are common (Lonsdorf & Merz, 2017), stable (Torrents-Rodas et al. 2014; Fredrikson, Annas, Georgiades, Hursti & Tersman, 1993; Zeidan et al. 2012) and have been associated with psychopathology (Duits et al. 2015; Nees, Heinrich & Flor, 2015; Lonsdorf & Merz 2017). In this study we examined the whole brain correlates of such differences in a large sample (*N* = 285) using simultaneous SCR and fMRI recordings. The fear conditioning task produced significantly larger SCRs to the CS+ than to the CS−, indicating successful conditioning at the group level. However, in line with previous research (Lonsdorf & Merz, 2017), there were also substantial individual differences in magnitude of conditioned response to the CS+ relative to the CS−. The SCR difference scores (CS+ - CS−) were calculated and correlated with whole brain activation to the CS+ relative to the CS− on a voxel-by-voxel basis. Results demonstrated that individuals showing heightened conditioned SCR to the CS+ relative the CS− also showed heightened neural activity to the CS+ relative to the CS− in dorsal anterior cingulate cortex (dACC)/anterior midcingulate cortex (aMCC), right anterior insula, bilateral temporoparietal junction/superior temporal gyri, right frontal operculum, bilateral dorsal premotor cortex (BA6), right superior parietal lobe/postcentral gyrus and a right-lateralized midbrain region in areas consistent with the periaqueductal gray and reticular formation. Furthermore, a ROI analysis indicated that responses in bilateral amygdala were correlated to conditioned SCR. Hierarchical regression analysis suggested, however, that cortical and midbrain regions contributed to individual differences in conditioned SCR beyond the amygdala. On the contrary, the amygdala did not show evidence of any unique contribution to SCR beyond the whole brain correlates. When examining the unique contributions of all neural correlates relative to each other, the right midbrain region showed a significant unique contribution, although this finding should be interpreted with caution since it did not replicate in a regression model excluding the amygdala. All regional activations demonstrated a stronger correlation with SCR to the CS+ compared to the CS−, although some correlations did not survive Bonferroni correction. This indicated that neural activity was primarily associated with heightened conditioned SCR to the CS+.

As expected, our whole brain analysis replicated previously reported correlations between individual differences in conditioned SCR and brain activity in the dACC and insula (Marin et al., 2019) as well as the amygdala (Labar et al 1998; Phelps et al 2004; Petrovic et al. 2008; MacNamara et al. 2015; Marin et al. 2019). We also identified correlations in novel regions consisting of bilateral temporoparietal junction/superior temporal gyri, right frontal operculum, bilateral dorsal premotor cortex, right superior parietal lobe and right midbrain. Notably, all whole brain correlates found in the present study belong to the set of regions consistently activated during human fear conditioning (Fullana et al., 2016). This indicates that these regions not only activate to the CS+ relative to the CS− in general, but that the magnitude of this activation also co-varies with individual differences in the magnitude of conditioned responding indexed by SCR. This is consistent with the proposal by Fullana et al. (2016; based on findings by e.g. Cameron, 2009; Craig, 2009; Critchley & Harrison, 2013; Medford & Critchley, 2010) that these regions, especially the dACC and anterior insula, are part of a large-scale neural network regulating autonomic responding. Reasonably, increased autonomic activation would correlate with larger responses in neural regions regulating autonomic processing.

Notably, various research groups other than Fullana et al. (2016) have examined the functional connectivity and network-structure between the regions reported in the present study (see e.g. Uddin, Yeo & Spreng, 2019, for a review). While Fullana et al. (2016) emphasize their role in autonomic-interoceptive processing and conscious awareness, others have proposed a more general function of detecting and preparing response to salient events across homeostatic, affective and cognitive domains (which includes, but also extend beyond, autonomic-interoceptive processing and autonomic regulation; see e.g. Seeley et al., 2007; Menon & Uddin 2010; Uddin 2015; Menon 2015; Uddin et al., 2019). We follow guidelines by Uddin et al. (2019) and refer to this network as the ‘midcingulo-insular network’. This name was chosen to reflect the anatomy and to avoid multiple functional labels dependent on context (e.g. ‘salience network’, Menon 2015; ‘cingulo-opercular network’, Dosenbach et al., 2008; ‘ventral attention network’, Corbetta, Patel & Shulman, 2008). Core regions of the midcingulo-insular network are the presently implicated dACC/aMCC (see Vogt, 2016, for a discussion regarding the naming of this region) and anterior insula, which are proposed to constitute the major input-output nodes within the midcingulo-insular network (Uddin et al., 2019; Menon, 2015). Specifically, the insula is thought to integrate cognitive, affective, interoceptive and homeostatic information while the dACC re-represents this summarized information in order to determine autonomic, behavioral and cognitive responding (Menon, 2015; Medford & Critchley, 2010). Efferent autonomic output from the dACC is proposed to be mediated by the periaqueductal gray (Menon, 2015) such as presently implicated (by overlap of the present midbrain activation cluster and coordinates previously denoted as the periaqueductal gray, see Linnman, Moulton, Barmettler, Becerra & Borsook, 2012). Notably, less well-characterized regions of the midcingulo-insular network also include the presently implicated bilateral temporoparietal junction/superior temporal gyri (Uddin et al., 2019; see e.g. Yeo et al. 2011; Seeley et al. 2007; Bzdok et al. 2013) and the presently implicated right frontal operculum (as part of what has previously been called the “cingulo-opercular network” or the “ventral attention network”; Uddin et al., 2019; see e.g. Corbetta et al. 2008). Right lateralized temporoparietal junction and frontal operculum appear to be specifically recruited during exogenous salience detection (Uddin et al., 2019), which was assumed in the current study. Finally, the presently implicated amygdala is also proposed to constitute a major subcortical node within this network (Menon 2015; Uddin et al., 2019). Thus, most of the regions implicated in the present analysis belong to this same well-replicated network and are consistent with its role in regulating autonomic responding. Exceptions to this proposal include bilateral dorsal premotor cortex. However, the premotor cortex is known to constitute a major cortical elicitor of the SCR (Dawson et al., 2007; Boucsein, 2012), making it a plausible link between midcingulo-insular activity and peripheral skin conductance. Consequently, based on our findings and previous theory we propose that individual differences in conditioned SCR may originate from altered activity within the midcingulo-insular network (Uddin et al., 2019) in conjunction with dorsal premotor cortex.

Within the midcingulo-insular network, the amygdala has been proposed to contribute to the cortical processing of salience through biasing cortical input based on context-specific negatively valenced cues (Menon et al., 2015). This is consistent with its role in learning, storing and responding to threat-conditioned cues, such as demonstrated in animal models (Ledoux 2000; Davis 2000). However, the midcingulo-insular network is also proposed to base salience processing on integrated cognitive information and subjective experience (Uddin et al. 2019; Menon 2015), such as processed in the anterior insula (see e g Medford & Critchley 2010 for a more detailed account), and such processes have been dissociated from the amygdala (e.g. Bechara et al., 1995; Tabbert, Stark, Kirsch, & Vaitl, 2006; for a discussion see e.g. LeDoux & Pine, 2016; LeDoux & Brown, 2017). This implies that amygdala activity should only have a partial influence on overall midcingulo-insular activity and consequent behavioral, autonomic and cognitive responding. The present finding that a number of midcingulo-insular regions explained significant individual variation in conditioned SCR beyond the amygdala is consistent with this model. Specifically, amygdala activity may serve as one among several inputs in determining the overall salience of a conditioned stimulus, while another important input is explicit cortical processing beyond the amygdala. Based on the overall input of salience information, integrated in the anterior insula, the strength of autonomic output may then be determined.

It has been suggested that understanding sources of individual differences in fear conditioning may uncover individual risk/resilience factors with respect to fear and anxiety that may ultimately aid the understanding and treatment of fear-related psychopathology (Lonsdorf & Merz, 2017). Our proposal that individual differences in conditioned SCR covary with activity in a large-scale neural network substantiating autonomic-interoceptive processing and salience detection may highlight such a source. Indeed, overactivity within the midcingulo-insular network has previously been suggested to underlie pathological anxiety (Menon, 2011; Menon 2015), as patients with anxiety disorders show altered functional connectivity within this network (Peterson, Thome, Frewen & Lanius 2014) as well as hyperactivity within the anterior insula (Paulus & Stein, 2006; Stein, Simmons, Feinstein & Paulus, 2007) and amygdala (Etkin & Wager, 2007) nodes of the network. Based on these and similar findings, it has been suggested that increases in anxiety and neuroticism may result from excessive processing of emotion-related salient cues (Menon, 2011) and/or alterations in interoceptive-autonomic processing (e.g. Paulus & Stein 2006; Medford & Critchley, 2010). As pathological fear and anxiety has also been associated with altered SCR discrimination during fear conditioning (Duits et al. 2015; Nees, Heinrich & Flor 2015) and as our results indicate that SCR discrimination covary with midcingulo-insular activity, our results are consistent with this anxiety model. Specifically, altered midcingulo-insular activity in response to threat-related cues may constitute a risk factor, biomarker or possibly even a cause of pathological fear and anxiety. We recommend future studies to continue examining this possibility, as it may have implications for the understanding and treatment of fear-related psychopathology.

One limitation of the present study is the use of social stimuli as CSs. While the general CS+>CS− BOLD contrast analyzed in the present study largely demonstrated a pattern of activation typical to fear conditioning (Fullana et al., 2016; see Supplementary Information 6), the potential influence of social threat processing cannot be entirely ruled out. Therefore, we recommend replicating our study using non-social stimuli as CSs.

In summary, the present study reports the largest to date study on individual differences in autonomic fear conditioning and brain activation. We are reporting several novel areas whose activation predict individual differences in conditioned SCR as well as replicating previous findings from smaller studies. Our results are consistent with the activation of a large-scale midcingulo-insular network substantiating autonomic-interoceptive processing and salience detection. We propose that altered activity within this network underlie individual differences in conditioned SCR and possibly autonomic regulation in pathological fear and anxiety. Future research should continue investigating this network as well as its possible relationship to fear and anxiety. Ultimately, such efforts may uncover the mechanisms of fear-related psychopathology and aid its treatment.

## 4. Materials and Methods

### 4.1 Subjects

This study was part of a twin study of genetic influences on fear related brain functions. Results describing genetic influences on SCR and fMRI responses during fear conditioning will be published elsewhere. 311 adult mono- and dizygotic twin volunteers were recruited from the Swedish Twin Registry (Svenska Tvillingregistret). Six participants were unable to participate in the fMRI-procedure and were excluded before data collection. After data collection, another twenty participants were excluded from analysis due to one or several of the following reasons: unsuccessful recording of skin conductance responses (2 participants); loss of brain imaging data due to excessive head-movement (5 participants); participant failure to comply with task instruction regarding button presses in at least 80% of trials (11 participants; see section 4.3.1); participant use of psychotropic medication (7 participants). Thus, 285 participants (female = 167, mean age = 33.92 years, *SD* = 10.11 years) were included in the analyses below. All participants passed the following exclusion criteria: pregnancy, inability to lie still for a 1 h duration, intolerance of tight confinements, ongoing psychological treatment, use of psychotropic medications, metal objects in the body (due to surgery, fragmentation etc.), current alcohol or drug related problems. However, in order to ensure the reliability of our findings we also performed an additional supplementary analysis where all participants with fMRI and SCR data were included (*N* = 303; see Supplementary Information 4). Participants provided written informed consent in accordance with guidelines from the Regional Ethics Review Board in Uppsala and received SEK 1000 as reimbursement for their participation. The study protocol was approved by the Regional Ethics Review Board in Uppsala.

### 4.2 Materials

#### 4.2.1 Stimuli and Contexts

For the fear conditioning task, two male three-dimensional virtual humanoid characters and a virtual environment (Figure 3) were created in Unity (version 5.2.3, Unity Technologies, San Francisco, CA). The virtual environment consisted of a room with four red colored brick walls, grey concrete roof and a wooden floor.

**Figure 3.**
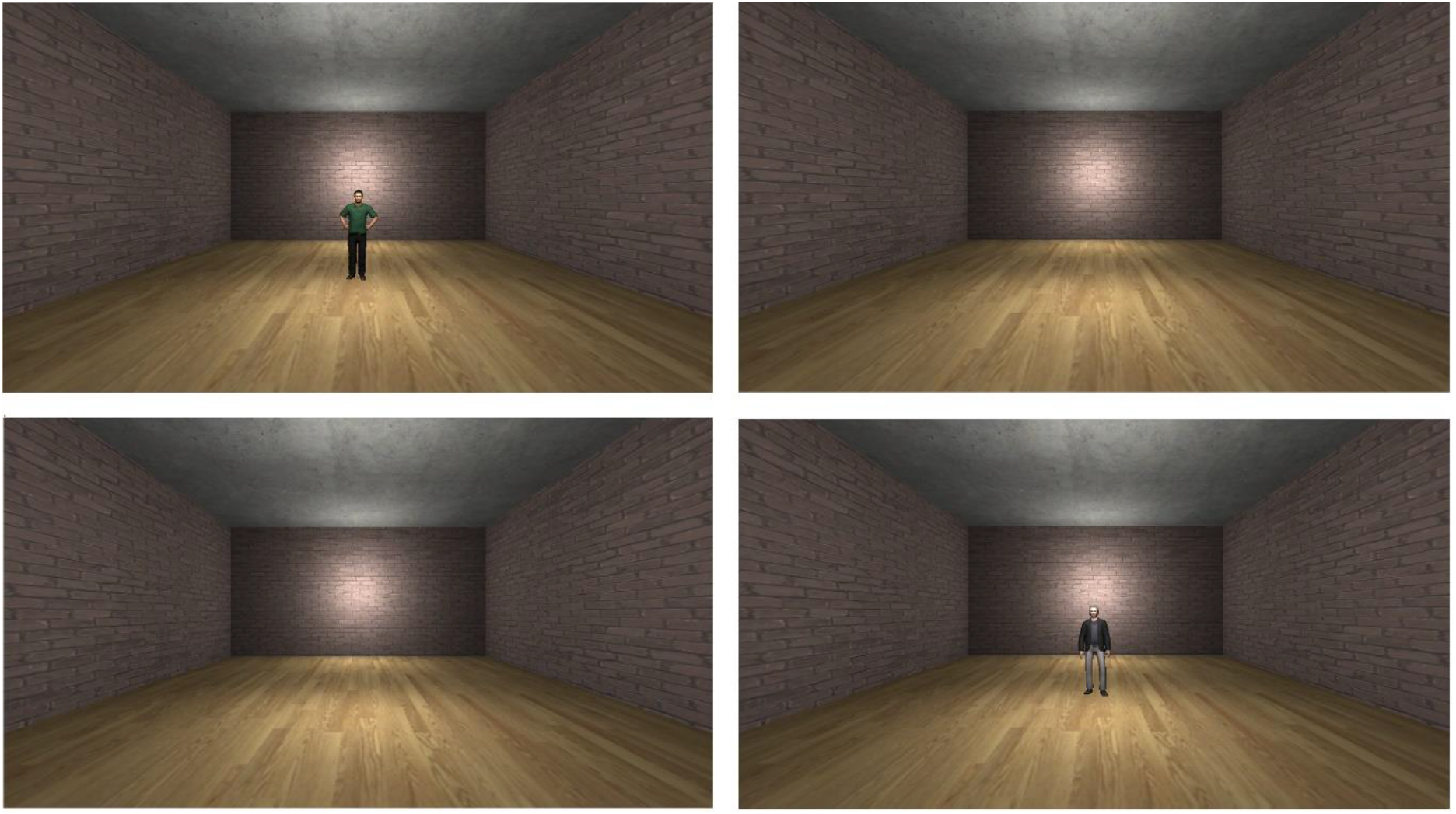
Experimental setup. Two male characters were displayed in the scanner during the fear conditioning task (top left, bottom right). One character predicted an electric shock (CS+) whereas the other served as a control stimulus and was never followed by shock (CS−). Between character presentations participants viewed the empty virtual environment (top right, bottom left).

#### 4.2.2 Stimulus Presentation Software

Contexts and stimuli were presented on a flat screen in the MR scanner with the help of a projector (Epson EX5260). The computer running the stimulus presentation used a custom version of Unity (version 5.2.3, Unity Technologies, San Francisco, CA) and communicated with BIOPAC (BIOPAC Systems, Goleta, CA) through a parallel port interface. The software for the parallel port interface was custom made and used standard .NET serial communication libraries by Microsoft (Microsoft Corporation, Albuquerque, New Mexico).

#### 4.2.3 Brain imaging

Imaging data were acquired using a 3.0 T scanner (Discovery MR750, GE Healthcare) and an 8-channel head-coil. Foam wedges, earplugs and headphones were used to reduce head motion and scanner noise. We acquired T1-weighted structural images with whole-head coverage, TR = 6400 ms, TE = 28 ms, acquisition time 6.04 min and flip angle 11 (degrees). Functional images were acquired using gradient echo-planar-imaging (EPI), TR = 2400 ms, TE = 28 ms, flip angle = 80 (degrees), slice thickness 3.0 mm with no spacing, axial orientation, frequency direction R/L, interleaved bottom up. Higher order shimming was performed and the number of dummy scans before the experiment was 5. Number of slices were 47 and voxel size 3.0 mm^3^.

#### 4.2.4 Skin conductance responses

Skin conductance recording was controlled with the MP-150 BIOPAC system (BIOPAC Systems, Goleta, CA). Radio-translucent disposable dry electrodes (EL509, BIOPAC Systems, Goleta, CA) were coated with isotonic gel (GEL101, BIOPAC Systems, Goleta, CA) and placed on the palmar surface of the left hand. The signal was high-pass filtered at 0.05Hz and SCRs were scored using Ledalab software package (Benedek & Kaernbach, 2010) implemented in Matlab (Mathworks, Inc., Natick, MA). SCR was analyzed using the maximum phasic driver amplitude 1-4 seconds after CS presentation for each participant.

Unlike previous studies considering the neural correlates of CS+>CS− SCR (Labar et al 1998; Phelps et al 2004; Petrovic et al. 2008; MacNamara et al. 2015; Marin et al. 2019), SCRs in the present study were range-corrected by Z transformation within individuals (Ben-Shakar, 1985). Z transformation increases sensitivity to conditioning related effects and prevents confounding by non-conditioning related individual differences in general SCR magnitude (Ben-Shakar, 1985). For comparison, however, neural correlations based on raw SCR scores were also examined and can be found in Supplementary Information 3.

Skin conductance was recorded without a low-pass filter. By using this recording procedure, we noticed that a small number of trials produced unreasonably high SCR values (e.g. 17 mS responding), likely due to electrode movement. As such extreme values may skew correlations even using standard scores, we excluded from all analysis trials with a raw value SCR score > 3 mS. This was based on previous research indicating a general maximum SCR between 2 and 3 mS using similar methodology as the one used in this paper (Boucsein, 2012). Using this criterion 97/12200 trials were excluded from analysis (0.795% of all trials).

### 4.3 Procedure

#### 4.3.1 Fear conditioning task

Two virtual characters served as CSs, one as a threat cue (CS+) predicting the US and the other as a control cue (CS−). CSs were presented on a screen in the MR scanner at a distance of 2.7 m in the virtual environment. The relatively long distance of 2.7 m was selected in order for the effect of conditioning on SCR not to be occluded by proximal threat effects on SCR, as was observed in two previous studies by Rosén et al. (Rosén, Kastrati & Åhs, 2017; Rosén, Kastrati, Reppling, Bergkvist & Åhs, 2019). Participants were told prior to the experiment that they could learn to predict the US but were not told which character served as the CS+. Participants were furthermore instructed to select either “yes” or “no” with the click of a button immediately following each CS presentation, indicating which character would be followed by the US. The inclusion of the button presses was to assure participants’ attention during the task and data from these responses were not subjected to further analysis. Which of the two characters that served as CS+ and CS− was counterbalanced across participants using four different stimulus presentation orders. Each CS presentation lasted for 6 s followed by an inter-stimulus interval of 8-12 s with no CS being present, but the context still being displayed.

A habituation phase preceded the fear conditioning, where each CS was presented four times without reinforcement for a total of eight CS presentations. This was followed by the fear conditioning phase, during which each CS type (CS+ and CS−) was presented 16 times. The experimental task thus consisted of 40 CS presentations in total, with 8 for habituation and 32 for conditioning. During conditioning, eight of the CS+ presentations co-terminated with a presentation of the US (50% partial reinforcement schedule; in accordance with guidelines for increasing sensitivity to inter-individual differences in fear conditioning, see Lonsdorf & Merz, 2017). Total time for the fear conditioning task was 9 min and 47 s.

The US consisted of an electric shock delivered to the subjects’ wrist via radio-translucent dry electrodes (EL509, BIOPAC Systems, Goleta, CA). Prior to the experiment, the shock was calibrated using an ascending staircase procedure where shock intensity is increased until rated by participants as “uncomfortable” but not “painful” (Åhs, Kragel, Zielinski, Brady, & LaBar, 2015). US duration was 16 ms and controlled using the STM100C module connected to the STM200 constant voltage stimulator (BIOPAC Systems, Goleta, CA).

#### 4.3.2 Analysis of SCR data

SCR Z scores were averaged separately across CS+ and CS− trials within each participant. A paired samples t test was performed comparing the average CS+ SCRs to the average CS− SCRs at a level of significance of *α* = .05 using JASP software (JASP Team (2020). JASP (Version 0.14.1) [Computer software]). This allowed us to determine whether the fear conditioning task was successful in evoking greater SCR to the CS+ than to the CS−. Secondly, in order to examine the correlations between conditioned SCR and fMRI responses during fear conditioning, an SCR difference score was calculated for each participant by subtracting the average SCR to CS+ presentations from the average SCR to CS− presentations. The distribution of SCR difference scores was examined to ensure the validity and sensitivity of neural regression analyses (see Supplementary Information 1 regarding methodology and results of SCR distribution analysis).

#### 4.3.3 Analysis of fMRI data

Analyses of fMRI data were performed using SPM12 (Welcome Department of Cognitive Neurology, University College, London). Preprocessing of images was performed using interleaved slice time correction, realignment and co-registration to acquired T1 images. Quality of images was controlled using MRIQC (mask validation matching; Esteban et al. 2017) and movement parameters. Images were normalized to Montreal Neurological Institute (MNI) standard space. 8 mm FHMW Gaussian smoothing was applied to ameliorate differences in inter-subject localization.

First-level analysis used event-related modeling including regressors for CS+ habituation trials, CS− habituation trials, CS+ acquisition trials, CS− acquisition trials and US delivery in a general linear model (GLM). Regressors mapped to the intervals of 6 s where each type of stimulus was displayed in the scanner and was convolved with the hemodynamic response function to predict the fMRI time course (for the brief US, the regressor still mapped to the 6 seconds following US delivery). Also, 6 movement parameter regressors, derived from the image realignment, and a mean value intercept regressor were included in the model. At the second level, we first examined the overall whole brain CS+>CS− BOLD contrast to assess conditioning related effects. Voxel-wise statistical significance was calculated using t tests implemented in the SPM12 software with an alpha level of *α* = .05 using family-wise error (FWE) correction. As results regarding this contrast is not central to the present study they are published in the Supplementary Information 6. However, it should be noted that these results were typical for fear conditioning studies in general (see e.g. Fullana et al. 2016). In the present analysis, we proceeded to examine regional activity within the whole brain CS+>CS− BOLD contrast that covaried with individual differences in the SCR difference score. This was done by entering each individuals previously obtained differential SCR score (CS+ minus CS−) as a second level between-subjects regressor of their average CS+>CS− BOLD activation, effectively correlating the CS+>CS− SCR difference with the CS+>CS− BOLD difference. Voxel-wise statistical significance was again calculated using *t*-tests implemented in the SPM12 software with an FWE corrected alpha level of *α* = .05. Notably, this provided a local maxima height threshold of *t* = 4.36 given our sample size (*N* = 285). This roughly corresponds to a correlation coefficient of *r* = .25 given the conversion formula *r* = sqrt (*t*^2 /(*t*^2 + DF)), thus corresponding to an effect size within the medium range according to guidelines by Cohen (Cohen 1988; Cohen 1992).

Secondly, we aimed to replicate previous findings of an association between individual differences in amygdala response and SCR using region-of-interest (ROI) analysis. The amygdala ROI was defined using the automated anatomical labeling (AAL) library (Maldjian, Laurienti, Kraft, & Burdette, 2003) and included both right and left amygdala. Analysis was performed the same way as the previous whole brain correlation only this time individual SCR scores were correlated exclusively to contrast values within the amygdala ROI, thus increasing sensitivity within this theoretically implicated region.

Third, in order to compare the independent contributions to explaining individual differences in conditioned SCR of clusters of voxels that showed significant correlations, we extracted eigenvariates from whole brain significant clusters and the amygdala ROI. Extracted eigenvariates for each cluster were then entered as regressors in a hierarchical regression implemented in the JASP software. The unique contribution to SCR of a specific neural region could then be calculated by adding all other regional eigenvariates to the null model and examining significant (*p* < .05) *r*-squared change as a result of including the specific neural region’s eigenvariates in the model.

Finally, in order to determine if obtained neural correlates to SCR difference scores was driven more by SCR to CS+ or CS−, we correlated eigenvariates extracted from implicated neural regions and average SCRs to the CS+ and CS− separately. In this analysis we used square root transformed raw value SCRs in order to obtain roughly normalized SCRs without confounding of CS+ and CS− response magnitude such as when using Z transformation. As the distributions of square root transformed raw value SCRs to the CS+ and CS− did not meet criteria for normality (by visual inspection of histograms, QQ-plot and Shapiro-Wilke’s test showing p < .001; see Supplementary Information 2), we used Spearman’s *Rho* instead of Pearson’s *r* correlations. To compare the difference between CS+ and CS− correlations we used the Steiger (1980) direct comparison of dependent correlation coefficients such as implemented in free automated software by Lee and Preacher (2013), as this is the most robust way of testing the difference between dependent Spearman coefficients (Myers & Sirois, 2006). To compensate for multiple testing during correlation comparisons, we used a Bonferroni corrected alpha level of *α* = .05/33 = .00151. This compensated for a total of 33 tests, reflecting 11 implicated neural regions each being correlated to CS+ and CS− separately, as well as each having these correlations compared in an additional test.

## Acknowledgements

This research was supported by grants from the Swedish Research Council (2014-01160, 2018-01322) and the Bank of Sweden Tercentenary Foundation (P20-0125).

