## Supplementary Information for "Whole Brain Correlates of Individual Differences in Skin Conductance Responses during Human Fear Conditioning"

### Supplementary Information 1. Distribution of individual SCR difference scores

*Method*

The distribution of SCR difference scores was examined using descriptive statistics, visual inspection of histogram, QQ-plot and box plot as well the Shapiro-Wilkes' test (Shapiro & Wilk 1965; Razali & Wah 2011), testing whether the obtained distribution deviated significantly from normality. These analyses were performed to ensure reasonable data with substantial individual differences in conditioned SCR, increasing the reliability and sensitivity of later regression analysis. Notice, however, that normality of the predictor variable is not an assumption in regression analysis using the general linear model (see e.g**.** Fox, 2015).

*Results*.

Shapiro-Wilkes' test ($p < .05$; Shapiro & Wilk 1965; Razali & Wah 2011) and a visual inspection of histogram, QQ-plot and box plot revealed a roughly symmetrical distribution (skewness = -0.02, SE = 0.14) that deviated from normality due to excessive negative kurtosis (kurtosis = -0.62, SE = 0.29; see Figure S1). This indicated substantial individual differences in conditioned SCR scores and increased sensitivity of regression analyses.

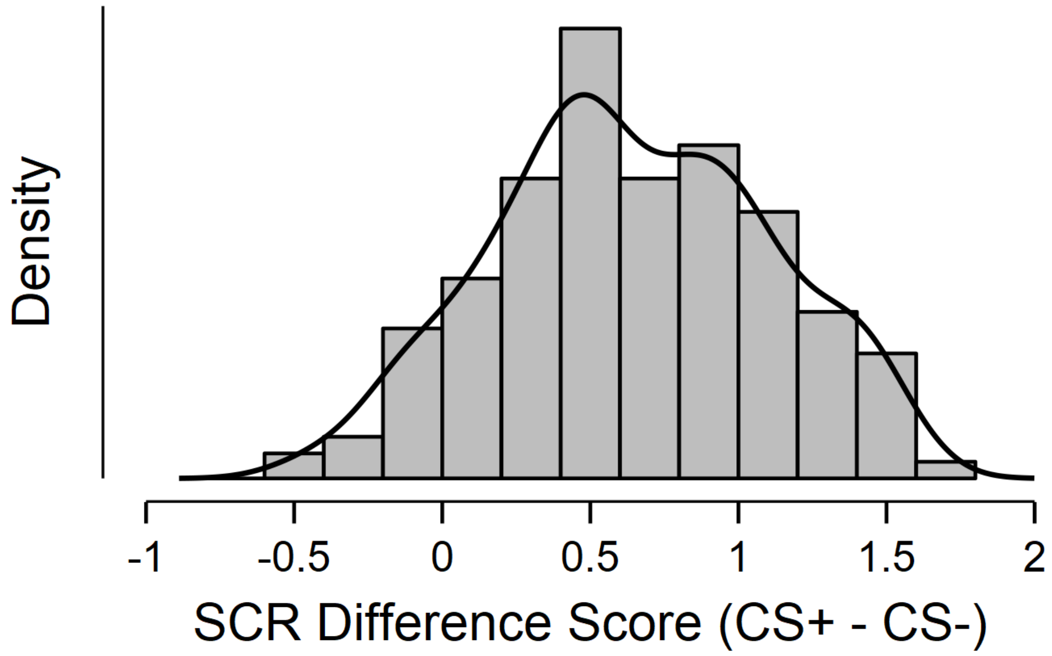


*Figure S1.* Histogram displaying distribution of SCR difference scores.

**Supplementary Information 2. Distribution of square-root transformed raw value SCRs to the CS+ and CS- separately.**

*Method*

Similar to the analysis of the distribution of individual differences in SCR difference scores above, we also examined the distribution of square root transformed raw value SCRs to the CS+ and CS- separately. This was done to determine the fit of using Spearman's *rho* correlations or Pearson's *r* correlations in the analyses considering CS+ and CS- responses separately (see Results section 2.2.3). Again, the distribution of SCRs was examined using descriptive statistics, visual inspection of histogram, QQ-plot and box plot as well the Shapiro-Wilkes' test (Shapiro & Wilk 1965; Razali & Wah 2011), testing whether the obtained distribution deviated significantly from normality.

*Results*

Shapiro-Wilkes' test ($p < .001$; Shapiro & Wilk 1965; Razali & Wah 2011) and a visual inspection of histogram, QQ-plot and box plot revealed non-normal distributions due to statistically significant positive skewness (right-tail) for both CS+ (skewness = 0.63, SE = 0.14; kurtosis = -0.11, SE = 0.29) and CS- (skewness = 0.84, SE = 0.14; kurtosis = 0.46, SE = 0.29). This indicated the need to use Spearman's *rho* correlations in the correlation analysis of these data.

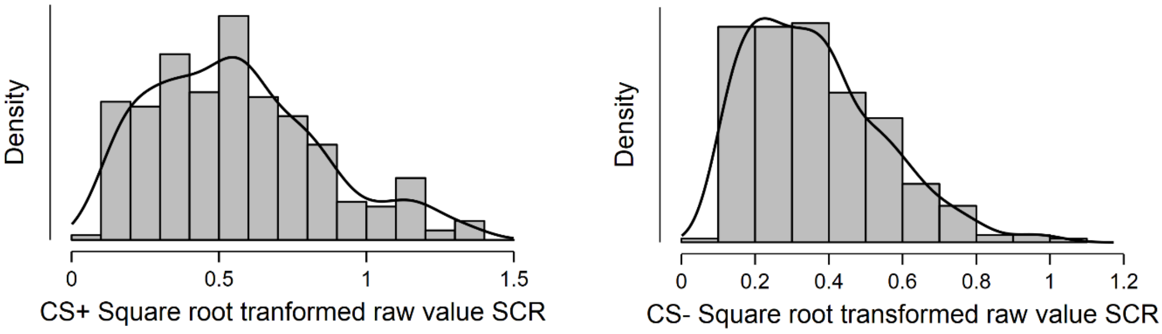


Figure S2. Histogram displaying distribution of average square root transformed raw value SCRs to the CS+ (left) and CS- (right).

**Supplementary Information 3. Whole brain correlation analysis using square-root transformed raw value SCR**

In order to examine if our choice of using range correction by Z transformation affected our results, we repeated our whole brain analysis using square-root transformed raw value SCR such as in previous analyses (e.g. MacNamara et al. 2015; Marin et al., 2019). Results demonstrated an almost identical pattern of activation to that in the main analysis of the main text, implicating right inferior frontal gyrus/frontal operculum (peak MNI coordinates: 56, 12, 2; cluster size: 144 voxels; $t = 5.66$), right temporoparietal junction (peak MNI coordinates: 64, -40, 20; cluster size: 116 voxels; $t = 5.64$), right superior frontal gyrus/dorsal premotor cortex (peak MNI coordinates: 16, 0, 66; cluster size: 125 voxels; $t = 5.54$), a right-lateralized midbrain region overlapping with the periaqueductal gray and reticular formation (peak MNI coordinates: 6, -30, -10; cluster size: 61 voxels; $t = 5.03$), dorsal anterior cingulate cortex (dACC)/anterior midcingulate cortex (aMCC; peak MNI coordinates: 4, 8, 40; cluster size: 21 voxels; $t = 4.58$), right anterior insula (peak MNI coordinates: 36, 22, 6; cluster size: 18 voxels; $t = 4.59$) and a small borderline significant activation in left temporoparietal junction (peak MNI coordinates: -62, -36, 22; cluster size: 6 voxels; $t = 4.49$). In comparison to the main results presented in Table 1 of the main text, this analysis thus implicated the same set of regions except not the left superior frontal gyrus/dorsal premotor cortex and right superior parietal lobe. Notably, this was two of the three smallest clusters of activation in the main analysis. Thus, our choice of using Z transformation had limited effect on our results.

Directly comparing SCR scores obtained using either Z transformation or square root transformed raw value SCRs demonstrated a high correlation between them (Spearman's rho $r_s = .86$; $p < .001$). Furthermore, Z transformed conditioned SCR scores were shown to correlate with raw response magnitude to both CS+ ($r_s = .62$; $p < .001$) and CS- ($r_s = .19$; $p = .001$), meaning that individuals who showed greater differentiation between CS+ and CS- also had overall higher magnitude SCR.

**Supplementary Information 4. Whole brain correlation analysis without participant exclusion (*N* = 303)**

In order to examine if our choice of participant exclusion in the main analysis affected our results, we repeated our whole brain analysis using the full sample of participants with both fMRI and SCR data. In addition to the 285 participants in the main analysis, this included an additional 5 participants with excessive head movement, an additional 11 participants that failed to comply with the task instruction regarding button presses in at least 80% of trials and an additional 7 participants that used psychotropic medication. Thus, this full sample size consisted of 303 participants.

Results using $N$ = 303 demonstrated an almost identical pattern of activation to that in the main analysis of the main text, implicating the dorsal anterior cingulate cortex/anterior midcingulate cortex (peak MNI coordinates: 6, 10, 38; cluster size: 101 voxels; $t$ = 5.06), right anterior insula (peak MNI coordinates: 36, 20, 6; cluster size: 16 voxels; $t$ = 4.54), right inferior frontal gyrus (peak MNI coordinates: 56, 10, 2; cluster size: 149 voxels; $t$ = 5.67), right temporoparietal junction (peak MNI coordinates: 64, -40, 20; cluster size: 91 voxels; $t$ = 5.82), right superior frontal gyrus/dorsal premotor cortex (peak MNI coordinates: 18, 0, 68; cluster size: 17 voxels; $t$ = 4.72), right midbrain in a region overlapping with the periaqueductal gray and/or superior colliculus (peak MNI coordinates: 10, -32, -10; cluster size: 74 voxels; $t$ = 5.29) and a small regional activation in left inferior frontal (peak MNI coordinates: -56, 0, 2; cluster size: 2 voxels; $t$ = 4.35). In comparison to the main results presented in Table 1 of the main text, this analysis thus implicated the same set of regions except the left superior frontal gyrus/dorsal premotor cortex, left temporoparietal junction and right superior parietal lobe. Notably, this was the three smallest clusters of activation in the main analysis. While this analysis using $N$ = 303 also implicated a new neural region, namely the left inferior frontal gyrus, this finding does not appear as robust as previous findings and should therefore be interpreted with caution.

In summary, both supplementary whole brain analyses (Supplementary Information 3 and 4) implicated largely the same set of regions as the main analysis of the main text (see Table 1). In particular, right-lateralized regional activations in the dorsal anterior cingulate cortex/anterior midcingulate cortex, anterior insula, inferior frontal gyrus, temporoparietal junction, superior frontal gyrus/dorsal premotor cortex and midbrain were consistent across analyses. Thus, findings regarding these regions appear robust. However, neural activations in the left superior frontal gyrus/dorsal premotor cortex, left temporoparietal junction, left inferior frontal gyrus and right superior parietal lobe were not consistently implicated. Hence, findings regarding these latter regions do not appear as robust. However, two important things should be noticed regarding these latter regions. First, our conclusions regarding the main findings of the study are not heavily dependent on the activation of these latter regions. Second, for reasons explained in the Materials and methods section of the main text, we consider the main analysis of the main text to be the most valid analysis.

**Supplementary Information 5. Beta coefficients from regression model predicting individual differences in conditioned SCR.**

**Table S1. Beta coefficients from regression model excluding amygdala ($H_0$) versus including amygdala ($H_1$)**

| Model | | Unstandardized | Standard Error | Standardized | t | p |
|---|---|---|---|---|---|---|
| $H_0$ | (Intercept) | 0.434 | 0.044 | | 9.773 | < .001 |
| | R IFG | 0.088 | 0.086 | 0.105 | 1.026 | 0.306 |
| | R TPJ | 0.136 | 0.088 | 0.133 | 1.551 | 0.122 |
| | R Midbrain | 0.178 | 0.107 | 0.122 | 1.671 | 0.096 |
| | R dPMC | 0.077 | 0.105 | 0.065 | 0.733 | 0.464 |
| | dACC | -0.064 | 0.092 | -0.072 | -0.699 | 0.485 |
| | R AI | 8.480e -4 | 0.080 | 9.570e -4 | 0.011 | 0.992 |
| | R SPL | 0.039 | 0.062 | 0.047 | 0.618 | 0.537 |
| | L SFG | 0.079 | 0.085 | 0.075 | 0.939 | 0.349 |
| | L TPJ | -0.011 | 0.070 | -0.013 | -0.162 | 0.872 |
| $H_1$ | (Intercept) | 0.413 | 0.046 | | 8.970 | < .001 |
| | Right Amygdala | -0.013 | 0.117 | -0.011 | -0.112 | 0.911 |
| | Left Amygdala | -0.159 | 0.119 | -0.136 | -1.334 | 0.183 |
| | R IFG | 0.105 | 0.087 | 0.125 | 1.203 | 0.230 |
| | R TPJ | 0.132 | 0.088 | 0.129 | 1.495 | 0.136 |
| | R Midbrain | 0.258 | 0.119 | 0.177 | 2.168 | 0.031 |
| | R dPMC | 0.047 | 0.106 | 0.039 | 0.437 | 0.662 |
| | dACC | -0.039 | 0.093 | -0.044 | -0.425 | 0.671 |
| | R AI | 0.003 | 0.080 | 0.004 | 0.039 | 0.969 |
| | R SPL | 0.051 | 0.063 | 0.063 | 0.822 | 0.412 |
| | L SFG | 0.101 | 0.086 | 0.096 | 1.180 | 0.239 |
| | L TPJ | 0.005 | 0.071 | 0.006 | 0.073 | 0.942 |

*Note.* Coefficient results obtained from regression analysis within the JASP software (JASP Team (2020). JASP (Version 0.14.1) [Computer software]) using eigenvariates from implicated whole brain regions as independent regressors of individual differences in conditioned SCR. Abbreviations: R = Right; L = Left; TPJ = Temporoparietal Junction; IFG = Inferior Frontal Gyrus; dACC = dorsal Anterior Cingulate Cortex; dPMC = dorsal Premotor Cortex; AI = Anterior Insula; SPL = Superior Parietal Lobe.

**Supplementary Information 6. Whole brain CS+>CS- BOLD contrast**

Examining the whole brain CS+>CS- BOLD contrast revealed a pattern of activation typical to fear conditioning studies in general (Figure S3; Table S2; for comparison see Fullana et al. 2016). A large major cluster (46343 voxels) was centered on bilateral insula spreading laterodorsally to bilateral inferior, middle, superior and precentral frontal gyri, medially to ventral striatum, caudally to thalamus and adjacent midbrain/brainstem regions (including the periaqueductal gray, reticular formation and dorsal pons), as well as spreading dorsally to medial wall cortex (including dACC, midcingulate cortex, pre-supplementary and supplementary motor areas and dorsal anterior precuneus). The cluster also encompassed large activation of bilateral middle and superior temporal gyri and supramarginal gyrus as well as inferior and superior parietal cortex. Also included in the cluster was bilateral amygdala activation as well as small portions of cerebellum. In separate clusters we observed activation in bilateral cuneus, bilateral middle prefrontal cortex and bilateral visual cortex.

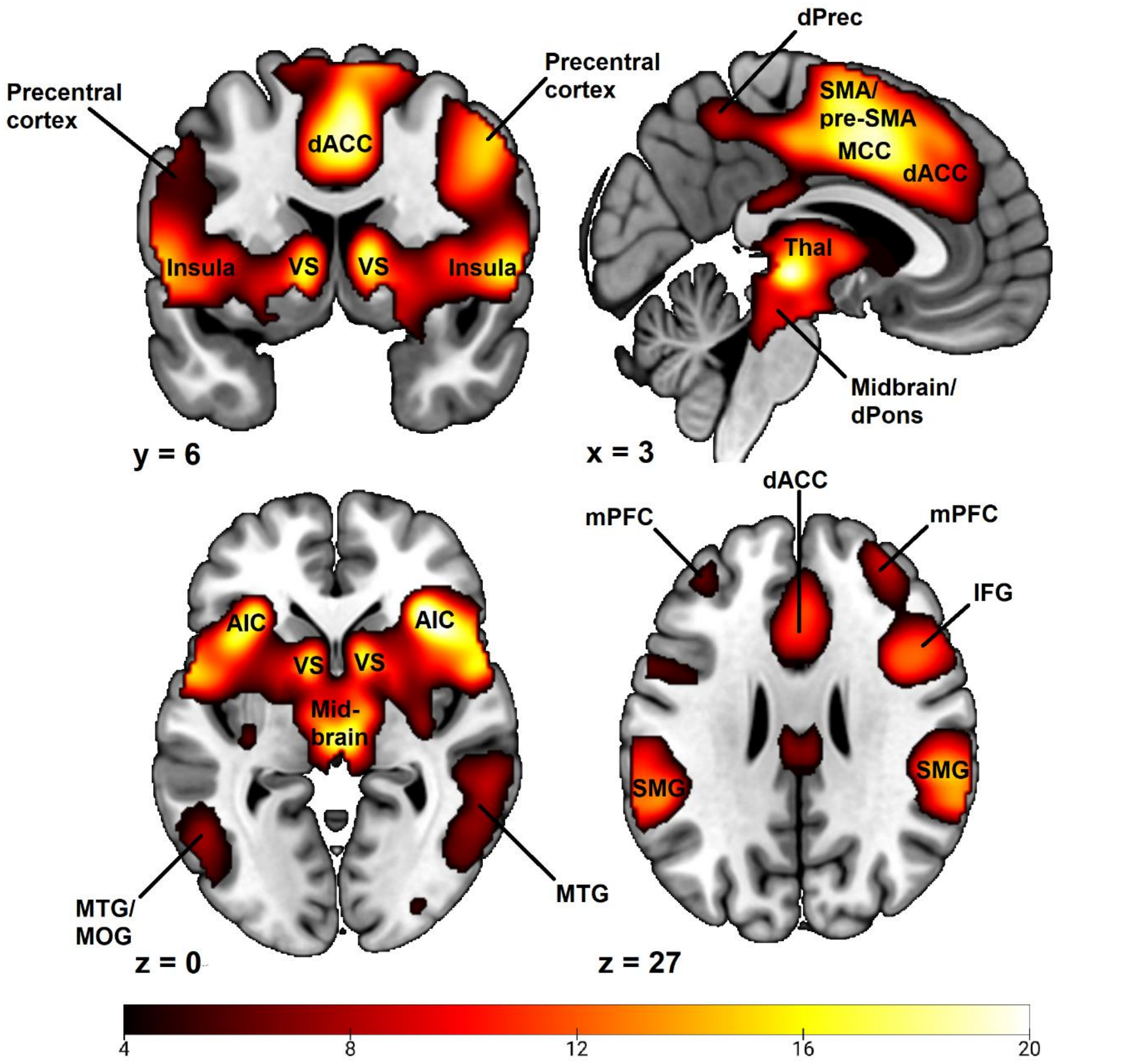


*Figure S3.* Whole brain CS+>CS- BOLD contrast activations. Color-coded *t* values ranging from $t = 4.0$ to $t = 20$. Statistical images thresholded at $p < 0.05$ FWE-corrected. Abbreviations: dACC, dorsal anterior cingulate cortex; VS, ventral striatum; SMA, supplementary motor area; Pre-SMA; pre-supplementary motor area; dPrec, dorsal precuneus; MCC, midcingulate cortex; Thal, thalamus; dPons, dorsal pons; AIC, anterior insula cortex; MTG, medial temporal gyrus; MOG, medial occipital gyrus; mPFC, middle prefrontal cortex; MFG, middle frontal gyrus; IFG, inferior frontal gyrus; SMG, supramarginal gyrus.

Table S2. Whole brain CS+ > CS- BOLD Contrast Activations

| | | | Peak MNI Coordinates | | |
|---|---|---|---|---|---|
| Cluster | Brain regions with peaks within cluster | *t* | x | y | z |
| **Cluster 1 (46343 voxels):** insula, ventral striatum, inferior, middle and superior frontal cortex, thalamus, midbrain/brainstem, anterior and midcingulate cortex, supplementary motor area, middle and superior temporal gyri, supramarginal gyrus, inferior and superior parietal cortex, amygdala, cerebellum. | - | >4.34 | - | - | - |
| | Right Insula | 21.71 | 36 | 24 | 2 |
| | Left Insula | 18.71 | -32 | 24 | -4 |
| | Left Insula | 17.55 | -40 | 18 | -4 |
| | Right Frontal Operculum | 17.06 | 56 | 6 | 2 |
| | Supplementary Motor Area | 19.24 | 4 | 6 | 50 |
| | Supplementary Motor Area/Midcingulate Cortex | 19.21 | 2 | 8 | 46 |
| | Right Caudate | 18.35 | 10 | 6 | 4 |
| | Left Caudate | 17.20 | -10 | 6 | 2 |
| | Thalamus | 20.10 | 4 | -24 | -2 |
| | Right Temporoparietal Junction | 17.86 | 50 | -32 | 20 |

| | | | | | |
|---|---|---|---|---|---|
| | Right Precentral Gyrus | 16.29 | 46 | 2 | 46 |
| **Cluster 2 (217 voxels):** Right Cuneus | | | | | |
| | Right Cuneus | 8.46 | 14 | -72 | 38 |
| **Cluster 3 (208 voxels):** Left Middle Prefrontal Cortex | | | | | |
| | Left Middle Prefrontal cortex | 6.33 | -38 | 46 | 24 |
| **Cluster 4 (69 voxels):** Left Cuneus, Left Posterior Precuneus | Left Posterior Precuneus | 6.04 | -10 | -74 | 38 |
| **Cluster 5 (61 voxels):** Left dorsal Cerebellum, Left Fusiform Gyrus | Left Cerebellum VI | 5.61 | -34 | -60 | -26 |
| | Left Cerebellum VI | 5.05 | -38 | -64 | -24 |
| | Left Fusiform Gyrus | 5.00 | -40 | -66 | -20 |
| | Left Cerebellum VI | 4.45 | -38 | -54 | -26 |
| **Cluster 6 (15 voxels):** Right Visual Cortex | Right Visual Cortex | 4.96 | 20 | -64 | 6 |
| **Cluster 7 (9 voxels):** Left Visual Cortex | Left Visual Cortex | 4.69 | -14 | -70 | 4 |
| **Cluster 8 (8 voxels):** Right Frontal Pole/Superior Orbital Gyrus | Right Frontal Pole | 4.56 | 26 | 60 | -6 |

*Note.* MNI coordinates and *t* values represent significant peak voxels within each cluster. Statistical significance was calculated using t tests with an FWE corrected alpha level of $\alpha = .05$ within the SPM software.
